# Invasive grass litter suppresses native species and promotes disease

**DOI:** 10.1101/2021.04.07.437244

**Authors:** Liliana Benitez, Amy E. Kendig, Ashish Adhikari, Keith Clay, Philip F. Harmon, Robert D. Holt, Erica M. Goss, S. Luke Flory

## Abstract

Plant litter can alter ecosystems and promote plant invasions by changing resource acquisition, depositing toxins, and transmitting microorganisms to living plants. Transmission of microorganisms from invasive litter to live plants may gain importance as invasive plants accumulate pathogens over time since introduction. It is unclear, however, if invasive plant litter affects native plant communities by promoting disease. *Microstegium vimineum* is an invasive grass that suppresses native populations, in part through litter production, and has accumulated leaf spot diseases since its introduction to the U.S. In a greenhouse experiment, we evaluated how *M. vimineum* litter and accumulated pathogens mediated resource competition with the native grass *Elymus virginicus*. Resource competition reduced biomass of both species and live *M. vimineum* increased disease incidence on the native species. *Microstegium vimineum* litter also promoted disease on the native species, suppressed establishment of both species, and reduced biomass of *M. vimineum*. Nonetheless, interference competition from litter had a stronger negative effect on the native species, increasing the relative abundance of *M. vimineum*. Altogether, invasive grass litter suppressed both species, ultimately favoring the invasive species in competition, and increased disease incidence on the native species.

## Introduction

Dead organisms and tissues can influence populations, communities, and ecosystems (Facelli and Pickett 1991, Renwick et al. 2007, Subalusky et al. 2017). Plant litter, for example, can release nutrients and toxins (Facelli and Pickett 1991), block light penetration to the soil (Molinari and D’Antonio 2020), mediate fire intensity (Flory et al. 2015), and host microorganisms that alter nutrient availability or disease (U’Ren and Arnold 2016). Litter can modify plant competition through chemical, physical, and biological processes. Specifically, litter can reverse the outcome of resource competition if the dominant competitor produces more litter and is more sensitive to litter than the inferior competitor (Kortessis et al. 2021). Litter impacts on resource availability (Farrer and Goldberg 2009, Eppinga et al. 2011, Aerts et al. 2017, Molinari and D’Antonio 2020) and fire regimes (D’Antonio and Vitousek 1992) can facilitate dominance of invasive plant species. However, research on invasive litter-mediated competition rarely considers pathogens that may reside in litter and infect living plants (Beckstead et al. 2012). Because invasive plants can accumulate pathogens over time (Goss et al. 2020), disease may increasingly influence litter-mediated competition with native species.

Invasive plants often dominate communities, produce abundant biomass (Vilà et al. 2011), and disproportionately contribute to litter, which can alter light, nutrients, soil moisture, and fire regimes (D’Antonio and Vitousek 1992, Farrer and Goldberg 2009, Wolkovich 2010, Flory et al. 2015). These processes can cause interference competition (Fig. 1a) and suppress establishment and growth of native (Walker and Vitousek 1991, Flory et al. 2015, Molinari and D’Antonio 2020) and invasive species (Chau et al. 2013, Warren et al. 2013). Conversely, litter can facilitate plant growth through increased soil moisture and nutrient availability (Wolkovich 2010, Chau et al. 2013). Regardless, litter effects must outweigh differences in resource competitive ability to affect the dominance of invasive plants (Kortessis et al. 2021).

**Figure 1.**
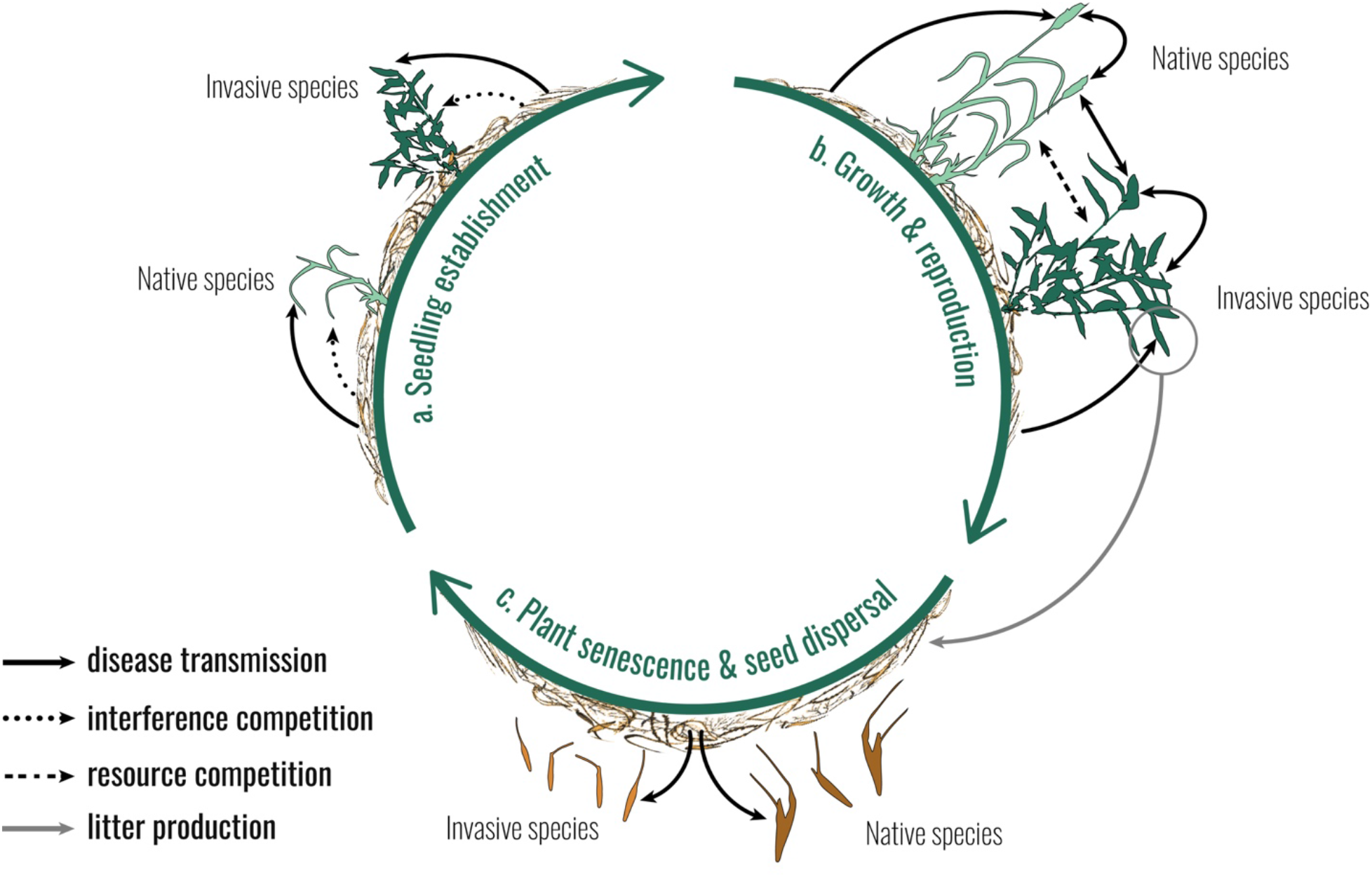
Litter is a potential source of pathogen propagules throughout the plant life cycle (a– c), which may be transmitted among live plants (b). Litter also has physical, chemical, and other biological effects on plant establishment and growth, resulting in interference competition (a). The net effects of litter can alter establishment (a), growth, reproduction (b), and seed survival (c). Litter may mediate invasive plant dominance (here, *M. vimineum*) over native species (here, *E. virginicus*) if the species differ in their sensitivity to negative effects of litter.

Litter can mediate wild plant diseases (Facelli et al. 1999, Whitaker et al. 2017), which has long been recognized in crop systems (Cook et al. 1978). Plant pathogens associated with litter (Chand et al. 2002) can transmit following plant dormancy (Fig. 1a) and throughout the plant life cycle (Fig. 1b-c; Beckstead et al. 2012). Litter may also indirectly promote infection by altering microenvironments (Bonanomi et al. 2011, Beckstead et al. 2012). Invasive litter impacts on disease are understudied, likely because invasive plants often escape native pathogens upon introduction, and pathogen accumulation takes time (Goss et al. 2020). However, after invasive plants have acquired pathogens in their new range, litter may take on the additional role of promoting disease. It is unclear whether litter’s impacts on disease, like its impacts on fire regimes and resource availability (Facelli and Pickett 1991), can mediate competition between native and invasive species.

Here, we experimentally investigated the potential for invasive plant litter containing accumulated pathogens to mediate competition with a native species. *Microstegium vimineum* (stiltgrass) is an invasive grass that has accumulated foliar fungal pathogens since its introduction (Stricker et al. 2016) and suppresses native species through resource and interference competition (Flory and Clay 2010, Flory et al. 2015). In a greenhouse experiment, we manipulated *M. vimineum* litter and competition between *M. vimineum* and the co-occurring native grass *Elymus virginicus* (Virginia wild rye; Flory et al. 2011). We hypothesized that litter would promote disease on and interfere with both species (Fig. 1). We expected that litter would reverse the competitive dominance of *M. vimineum* if it is more sensitive than *E. virginicus* to litter-mediated disease and interference competition (Kortessis et al. 2021).

## Methods

### Study System

*Microstegium vimineum* (Trin.) A. Camus is a C-4 annual grass that is invasive in the eastern U.S. (USDA 2020). It grows rapidly, produces large quantities of litter, and reduces plant abundances (Flory and Clay 2010, Flory et al. 2017), possibly through increased fire intensity (Flory et al. 2015), physical interference (Flory and Clay 2010), or toxins (Pisula and Meiners 2010). Its litter also may harbor pathogens (Chand et al. 2002). Over the past 20 years, leaf spot diseases caused by *Bipolaris* fungal pathogens that can reduce *M. vimineum* biomass and seed production have been identified in U.S. *M. vimineum* populations (Flory et al. 2011, Stricker et al. 2016). *Bipolaris* pathogens also infect the native perennial grass *E. virginicus* (Flory et al. 2011, Lane et al. 2020) and can reduce its biomass production (Kendig et al. 2021).

### Experimental design

To evaluate the potential for invasive plant litter to mediate competition with a native species, we manipulated invasive *M. vimineum* litter amount and plant species composition in a greenhouse experiment. In April 2018, we identified a forested site heavily invaded by *M. vimineum* at Big Oaks National Wildlife Refuge (BONWR) in Madison, IN, USA (38.965630, - 85.364500). Dead *M. vimineum* leaves at the site showed symptoms of *Bipolaris* infection, including leaf spots with a dark brown border and light interior (Lane et al. 2020). Live *M. vimineum* and *E. virginicus* at the site had symptoms previously shown to be caused by *B. gigantea* and other *Bipolaris* spp. (Stricker et al. 2016, Lane et al. 2020).

In May 2018, we collected litter from the BONWR site and transported it to the University of Florida in Gainesville, FL, USA. To confirm the presence of *Bipolaris* spores, we first incubated 65 g of litter with 50 ml sterile deionized water at 27°C for 48 hours. We then washed the litter in 120 ml sterile deionized water with 0.1% Tween 20 (Sigma-Aldrich, St. Louis, MO, USA) and filtered rinsate through cheese cloth, resulting in a crude suspension of conidia. Conidia were quantified using a Bright-Line hemocytometer, where elongated multi-celled conidia with longitudinal cell walls characteristic of *Bipolaris* species were counted (Lane et al. 2020), yielding 5500 conidia/g in the litter (USDA-APHIS-PPQ permit PP526P-18-01688).

*Microstegium vimineum* has a mixed mating system of cleistogamous (obligately selfed, within leaf sheaths) and chasmogamous (potentially outcrossed, open) seeds (Baker and Dyer 2011). To minimize germination of cleistogamous seeds, we hand-picked them along with any non-*M. vimineum* material from the litter. We created three litter treatments within the range of litter observed at BONWR (Appendix S1): 0.91 g/pot (50 g/m^2^, “low”), 1.82 g/pot (100 g/m^2^, “medium”), 3.64 g/pot (200 g/m^2^, “high”), and a control treatment without litter (“none”). To enhance sporulation of pathogenic fungi in the litter, we incubated the litter for 56 hours at room temperature in one-gallon plastic bags with a paper towel saturated with deionized water.

We collected *M. vimineum* seeds from BONWR in fall 2015 and purchased *E. virginicus* seeds from Prairie Moon Nursery (Winona, MN, USA) in spring 2018. On June 15, 2018, we planted three treatments: 50 seeds of *E. virginicus*, 50 seeds of *M. vimineum* (both “alone”; i.e., intraspecific competition), and 50 seeds of each (“in competition”; i.e., inter- and intraspecific competition), into 1 L (15.2 cm diameter) plastic pots filled with Metromix 930 growing medium (Sungro Horticulture, Agawam, MA, USA) saturated with tap water. We added the litter treatments to pots, creating twelve treatments (four litter treatments crossed with three planting treatments) with six replicates each. The following day, we topped each pot with a clear plastic sheet (50.8 cm width, 17.8 cm height; 0.005 Grafix Dura-Lar^®^ film, Maple Heights, OH) formed into a cone to increase humidity and potentially enhance fungal sporulation. Cones were opened to cylinders as plants outgrew them. We placed pots under shade tents in the greenhouse to reduce heat stress (photosynthetically active radiation: 224 μMolm^−2^s^−1^ +/− 18 μMol m^−2^s^−1^), haphazardly rearranged them across two greenhouse benches weekly, and watered them daily.

### Data collection

To measure establishment, we counted the number of plants per pot, and to measure disease incidence, we counted the proportion of plants per pot with at least two foliar lesions. We measured establishment and disease incidence weekly for six weeks. Some *M. vimineum* seedlings grew in *E. virginicus*-only pots, likely from cleistogamous seeds left in the litter despite our attempts to remove them. We counted and removed these seedlings. After 10 weeks, we cut plant stems at the soil surface and stored each species from each pot in a separate paper bag at 4°C. We then counted the number of *M. vimineum* plants and *E. virginicus* tillers (individuals were not identifiable) per pot, and the proportion with two or more foliar lesions. Because *M. vimineum* had few lesions, we searched for a maximum of ten minutes per *M. vimineum* per pot. Plants were then oven-dried at 60°C to a constant mass and weighed.

To identify foliar pathogens, we selected five *M. vimineum* and six *E. virginicus* leaves with relatively high numbers of lesions or large lesions from seven treatments (*M. vimineum* planted with medium and high litter; *E. virginicus* planted with no, low, and high litter; and both species planted with low and high litter). We placed each leaf in a petri dish with 7 cm filter paper wetted with deionized water and incubated them at 26°C under 12h light/dark cycle for 24 hours. Under a dissecting microscope, we searched leaves for conidiophores and conidia, which we transferred to a V8 media agar plate using a sterile dissecting needle. We incubated plates at 26°C under 12h light/dark cycle for 5–7 days and identified the fungal genus or species based on conidia size and morphology (Lane 2020).

### Data analysis

To evaluate the effects of litter and interspecific competition on establishment and disease incidence, we fit generalized linear regressions with binomial error distributions (logit link) to the corresponding proportions for each species with litter mass, planting treatment, and their interaction as independent variables. We used maximum establishment observed across the greatest number of treatments (26- and 39-days post planting for *M. vimineum* and *E. virginicus*, respectively) and disease incidence collected post-harvest. To account for *M. vimineum* establishment from cleistogamous seeds, we averaged the number of *M. vimineum* seedlings in *E. virginicus*-only pots by litter treatment and added this value to the number of *M. vimineum* seeds planted in *M. vimineum*-only or “in competition” pots with corresponding litter treatments. In two cases, *M. vimineum* plants exceeded the sum of the number planted and estimated cleistogamous seeds so we increased the summed number to match the number of plants. To evaluate the effects of treatments on cleistogamous seed germination, we fit a generalized linear regression with a Poisson error distribution (logit link) to the number of *M. vimineum* plants per pot with the independent variables described above.

To evaluate the effects of litter and interspecific competition on average plant biomass, we log-transformed biomass divided by number of plants per pot for each species and fit normal linear regressions with the independent variables described above. To evaluate the effect of disease on establishment and plant biomass, we fit normal linear regressions to the residuals of the establishment and biomass regressions with disease incidence as the independent variable. To evaluate the effect of litter on relative abundance, we fit a normal linear regression to log-transformed *M. vimineum* biomass from “in competition” pots with litter as an independent variable and an offset variable of log-transformed total biomass, effectively creating a dependent variable of the log ratio of *M. vimineum* biomass to total biomass (Zuur et al. 2009).

Multiple response variables did not change monotonically with litter mass, so we refit all models with litter as a categorical variable (i.e., none, low, medium, high) and compared them to models with the continuous litter variable using Akaike information criterion (AIC). We present results from models with lower AIC values (see Appendix S2 for ΔAIC). We used backwards stepwise selection to test the statistical significance of interactions and, if applicable, main effects (Crawley 2007), comparing nested models with Chi-squared tests (binomial/Poisson) or F tests (normal). We performed analyses in R version 4.0.1 (R Core Team 2020).

## Results

An average of 86% of *M. vimineum* seeds established without litter and interspecific competition (Fig. 2a). The maximum litter amount reduced *M. vimineum* establishment 14.6% when it was alone and 7.0% when *E. virginicus* was present (litter–competition interaction: *P* = 0.04, Appendix S2: Table S1). However, more cleistogamous seeds in pots with more litter led to consistent *M. vimineum* density across litter amounts (Appendix S2: Table S2). *Elymus virginicus* establishment without litter and interspecific competition was 84% (Fig. 2b). The presence of litter reduced *E. virginicus* establishment 8.9% (*P* < 0.001), on average, but there was no significant effect of interspecific competition (*P* = 0.2, Appendix S2: Table S3).

**Figure 2.**
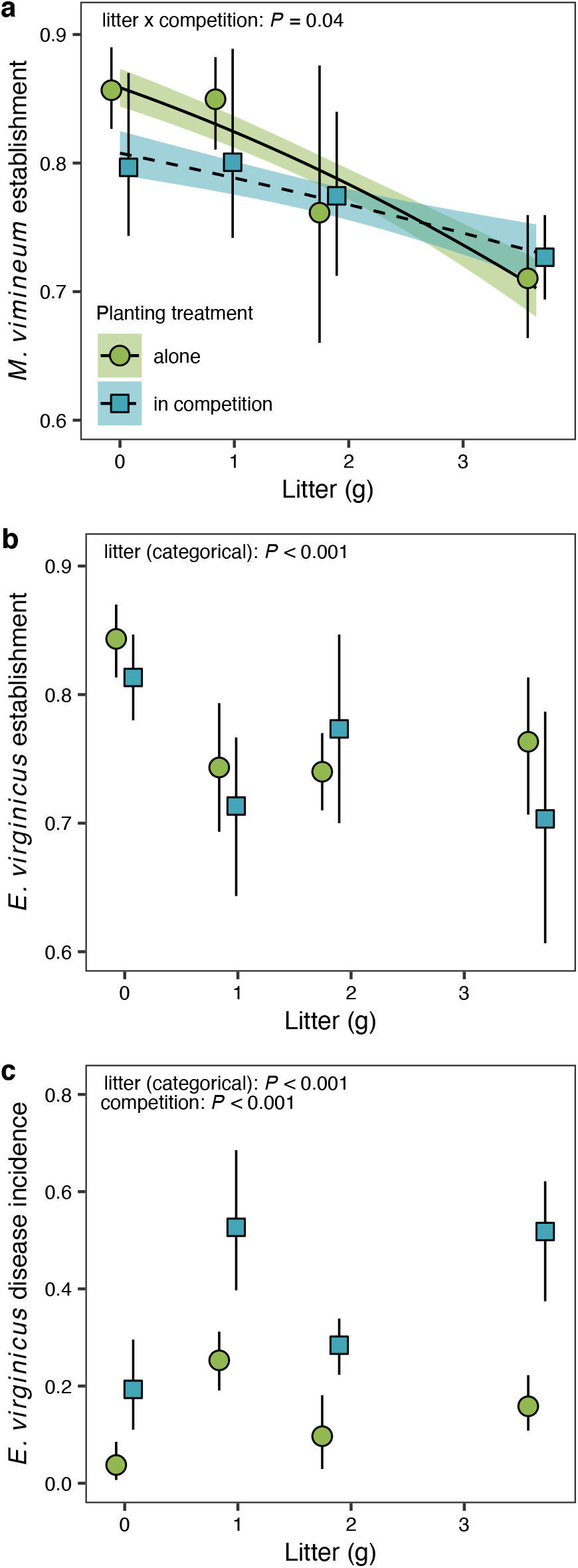
*Microstegium vimineum* litter reduced establishment of (a) *M. vimineum* and (b) *E. virginicus* and (c) increased disease incidence on *E. virginicus*. The effect of litter mass on (a) *M. vimineum* establishment depended on whether it was planted alone or with *E. virginicus* (lines and shading: model mean ± 1 SE). Presence of live *M. viminum* (c) increased disease incidence, (b) but did not affect *E. virginicus* establishment. Points and error bars (mean ± 95% CI) are nudged horizontally to reduce overlap.

*Microstegium vimineum* developed few lesions, with disease incidence of only 2% across all treatments (data not shown). *Elymus virginicus* had disease incidence of 3.8% without litter and interspecific competition (Fig. 2c). The presence of litter and interspecific competition increased *E. virginicus* disease incidence an average of 19.1% (*P* < 0.001) and 24.4% (*P* < 0.001), respectively. The interaction between litter and interspecific competition was not significant (*P* = 0.08, Appendix S2: Table S4). From the five *M. vimineum* and six *E. virginicus* leaves chosen for pathogen assessments, we identified *B. gigantea* on four *M. vimineum* leaves and another *Bipolaris* species on the fifth *M. vimineum* leaf and two *E. virginicus* leaves. We identified a *Curvularia* species, a common saprophyte, either alone or in combination with *Bipolaris* on one *M. vimineum* leaf and two *E. virginicus* leaves. We identified a *Cladosporium* species, a common endophyte, on a *M. vimineum* leaf that was also infected with *Bipolaris*.

*Microstegium vimineum* weighed an average of 0.9 g per plant without litter and interspecific competition (Fig. 3a). The maximum litter amount and interspecific competition reduced *M. vimineum* plant biomass 16.3% (*P* = 0.02) and 11.5% (*P* = 0.04), respectively. *Elymus virginicus* weighed an average of 0.2 g per plant without litter and interspecific competition (Fig. 3b). Litter did not significantly affect *E. virginicus* plant biomass (*P* = 0.1), but interspecific competition reduced it 84.6% (*P* < 0.001). Litter–competition interactions were not significant for either species (Appendix S2: Tables S5–S6). Disease incidence did not significantly affect *E. virginicus* establishment or plant biomass after accounting for treatment effects (Appendix S2: Tables S7–S8). *Microstegium vimineum* comprised 96.4% of the biomass in pots with both species and no litter (Fig. 3c). Litter increased the relative abundance of *M. vimineum* (*P* = 0.02), where the minimum litter amount increased it 1.8% and the maximum amount increased it 0.7% (Appendix S2: Table S9).

**Figure 3.**
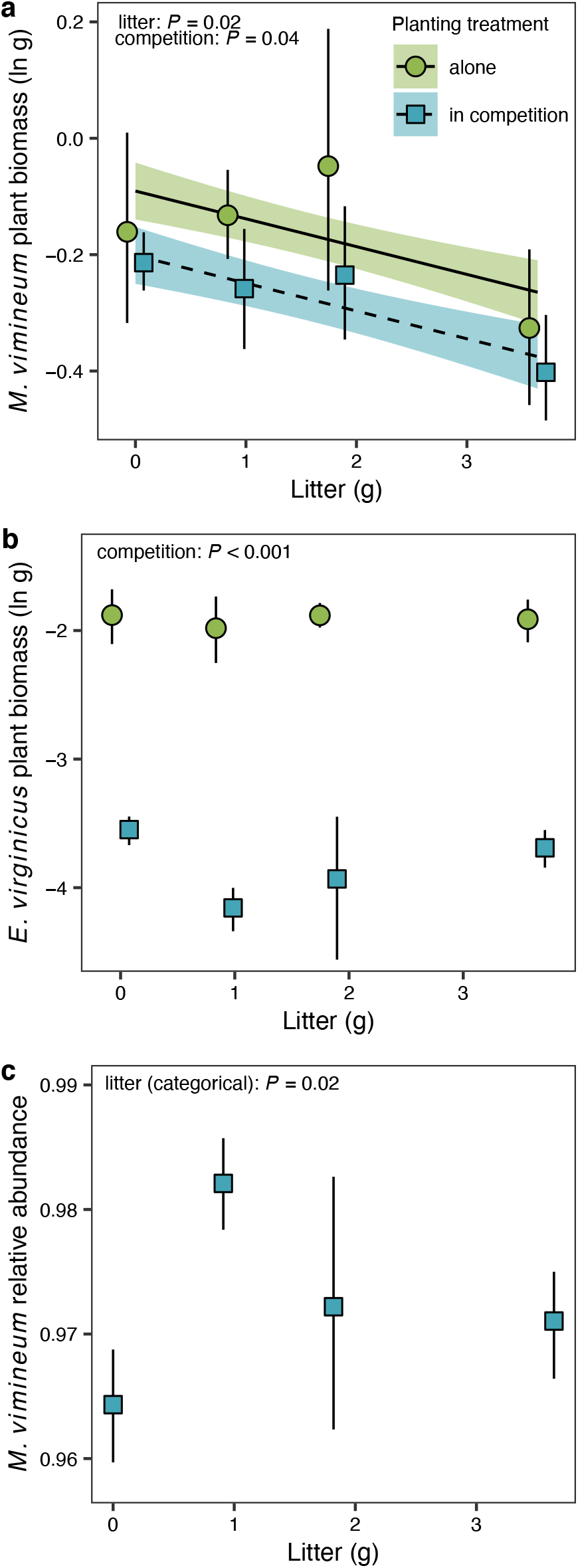
*Microstegium vimineum* litter reduced (a) *M. vimineum* plant biomass (lines and shading: model mean ± 1 SE) and increased (c) *M. vimineum* relative abundance. Competition reduced (a) *M. vimineum* and (b) *E. virginicus* plant biomass. Points and error bars (mean ± 95% CI) are nudged horizontally to reduce overlap.

## Discussion

Invasive plant litter can play an important role in community dynamics (Facelli and Pickett 1991, D’Antonio and Vitousek 1992, Eppinga et al. 2011) but the degree to which it influences communities through disease is unclear (Beckstead et al. 2012). We evaluated the impacts of litter from the invasive grass *M. vimineum* on disease incidence and competition with the native grass *E. virginicus*. Although litter negatively impacted both species, it ultimately increased the relative abundance of *M. vimineum*. Because disease incidence on *E. virginicus* did not significantly affect its establishment or biomass, it is likely that *M. vimineum* litter increased the relative abundance of *M. vimineum* through asymmetric interference competition and not disease. Therefore, invasive plant litter may drive interference competition and promote disease on native species, ultimately favoring the invasive species.

We expected pathogen transmission from litter to live plants to alter resource competition between the native and invasive species because pathogen exposure can reduce growth of both species (Stricker et al. 2016, Kendig et al. 2021). However, litter only promoted disease on *E. virginicus* and there were no significant effects of disease on *E. virginicus* establishment and biomass after accounting for treatment effects. Further, there were no significant interactive effects of litter and competition on biomass. Dry conditions, despite our use of humidity chambers, may have limited lesion formation on *M. vimineum* (Appendix S3) and disease severity on *E. virginicus*. Additionally, *M. vimineum* may have shed infected leaves between assessments (Vloutoglou and Kalogerakis 2000). However, live *M. vimineum* promoted disease incidence on *E. virginicus* and we hypothesize that *M. vimineum* biomass altered the microclimate to favor fungal sporulation. Therefore, *M. vimineum* may promote disease on co-occurring native species through both litter and live biomass, which could be problematic for more susceptible native species or environmental conditions conducive to greater disease severity. Lesions on *E. virginicus* plants without litter suggest an additional pathogen source, such as unsterilized soil or seeds. While we cannot conclude that infection came from litter, we isolated *Bipolaris* species from both litter and leaf spot lesions on both hosts.

Litter reduced *M. vimineum* establishment and biomass. This negative feedback between *M. vimineum* growth and litter production may regulate or reduce *M. vimineum* density over time (Tilman and Wedin 1991, Flory et al. 2017). While increasing litter amounts decreased *M. vimineum* establishment and biomass, *E. virginicus* was most sensitive to low amounts of litter. Stronger interference competition on *E. virginicus* than *M. vimineum* likely drove increased *M. vimineum* relative abundance, which was greatest with low litter. Because *M. vimineum* better tolerates low amounts of litter than *E. virginicus*, litter reinforces the negative impacts of *M. vimineum* on *E. virginicus* through resource competition (Kortessis et al. 2021). We did not isolate the mechanism by which *M. vimineum* litter suppressed plant establishment and growth, but *M. vimineum* litter reduces light penetration to the soil surface (Flory and Clay 2010) and the leaves contain phytotoxins (Pisula and Meiners 2010). Because litter reduced *M. vimineum* establishment, but deposited cleistogamous seeds, it may increase the proportion of *M. vimineum* germinating from cleistogamous seeds (obligate selfing) relative to chasmogamous seeds (potentially outcrossing), which could reduce genetic diversity of *M. vimineum* populations (Baker and Dyer 2011). To our knowledge, the role of litter in ecology and evolution of plant mating systems has not been studied and may be a fruitful direction of inquiry.

While the importance of invasive plant litter for resource availability (Farrer and Goldberg 2009, Aerts et al. 2017) and fire regimes (D’Antonio and Vitousek 1992) has been recognized for multiple systems, its role in mediating species interactions through disease has received little attention (Beckstead et al. 2012). We found that litter suppressed plant establishment and growth and promoted disease on the native species. While disease did not affect relative abundance, greater disease severity (e.g., due to more favorable environmental conditions) could affect *M. vimineum* competition with native species (Stricker et al. 2016). These processes can co-occur with interactions among live plants (Fig. 1), and here, live invasive grass reduced biomass and promoted disease on the native species. Thus, invasive plant impact assessments should not only consider effects of live plants on native communities and ecosystems but should also evaluate litter effects and their potential role in promoting disease.

## Acknowledgements

We thank Zobia Chanda, David Notman, Penny Reif, and Callie San Antonio for data collection assistance, Christopher Wojan, Briana Whitaker, Brett Lane, and Michael Barfield for advice on experimental design and analyses, Joe Robb for research access to BONWR, and Wesley Lewis for creating Fig. 1. This work was funded by USDA award number 2017-67013-26870 as part of the joint USDA- NSF-NIH Ecology and Evolution of Infectious Diseases program.

## Notes

### Competing Interest Statement

The authors have declared no competing interest.

