## Appendix S1 for "Invasive grass litter suppresses native species and promotes disease"

**Appendix S1** for "Invasive grass litter suppresses native species and promotes disease " by Liliana Benitez, Amy E. Kendig, Ashish Adhikari, Keith Clay, Philip F. Harmon, Robert D. Holt, Erica M. Goss, and S. Luke Flory

On April 4, 2018, we collected litter from six haphazardly selected 0.25 $m^2$ plots at the BONWR site described in the main text. We cut the litter around the inside edges of the PVC used to delineate the plots and put it in plastic bags. The following week, it was weighed at the University of Indiana (Bloomington, IN, USA). We did not separate *M. vimineum* litter from other types of litter, but *M. vimineum* appeared to be the dominant species.

**Table S1.** Weights of litter harvested from BONWR.

| ID | Litter density (g/0.25 $m^2$) | Litter density (g/$m^2$) |
|---|---:|---:|
| A | 49.34 | 197.36 |
| B | 72.75 | 291.00 |
| C | 92.44 | 369.76 |
| D | 91.19 | 364.76 |
| E | 114.72 | 458.88 |
| F | 51.18 | 204.72 |
| Average | 78.60 | 314.41 |
| Standard error | 10.48 | 41.93 |
