## Appendix S2 for "Invasive grass litter suppresses native species and promotes disease"

**Appendix S2** for "Invasive grass litter suppresses native species and promotes disease " by Liliana Benitez, Amy E. Kendig, Ashish Adhikari, Keith Clay, Philip F. Harmon, Robert D. Holt, Erica M. Goss, and S. Luke Flory

**Table S1.** Model summary of *M. vimineum* establishment (proportion of seeds, binomial distribution, logit link). Estimates, standard errors (SE), z statistic, and associated *P*-value from maximal model (*n* = 48). Deviance and *P*-value from Chi-squared tests of nested models. Model comparison with categorical litter model: ΔAIC = 4.7.

| Predictor | Estimate | SE | z | Pr(>\|z\|) | Deviance | Pr(>Chi) |
|---|---|---|---|---|---|---|
| Intercept | 1.80 | 0.12 | 14.98 | < 0.001 | | |
| Litter (g) | -0.26 | 0.05 | -5.22 | < 0.001 | | |
| *Elymus* present | -0.37 | 0.16 | -2.26 | 0.02 | | |
| Litter:*Elymus* present | 0.14 | 0.07 | 2.01 | 0.04 | -4.06 | 0.04 |

**Table S2.** Model summary of *M. vimineum* density (plants per pot, Poisson distribution, logit link). Estimates, standard errors (SE), z statistic, and associated *P*-value from maximal model (*n* = 48). Predictor variables without estimates were omitted during model selection (deviances and *P*-values from Chi-squared tests of nested models). Model comparison with categorical litter model: ΔAIC = 5.3.

| Predictor | Estimate | SE | z | Pr(>\|z\|) | Deviance | Pr(>Chi) |
|---|---|---|---|---|---|---|
| Intercept | 3.74 | 0.05 | 76.82 | < 0.001 | | |
| Litter (g) | -0.001 | 0.02 | -0.03 | 0.98 | -0.62 | 0.43 |
| *Elymus* present | -0.07 | 0.07 | -0.94 | 0.35 | -0.22 | 0.64 |
| Litter:*Elymus* present | 0.03 | 0.03 | 0.83 | 0.41 | -0.69 | 0.41 |

**Table S3.** Model summary of *E. virginicus* establishment (proportion of seeds, binomial distribution, logit link). Estimates, standard errors (SE), z statistic, and associated *P*-value from maximal model ($n$ = 48). Predictor variables without estimates were omitted during model selection (deviances and *P*-values from Chi-squared tests of nested models). Model comparison with continuous litter model: ΔAIC = 8.4.

| Predictor | Estimate | SE | z | Pr(>\|z\|) | Deviance | Pr(>Chi) |
|---|---|---|---|---|---|---|
| (Intercept) | 1.68 | 0.16 | 10.60 | < 0.001 | | |
| Low litter | -0.62 | 0.21 | -3.00 | 0.003 | | |
| Medium litter | -0.64 | 0.21 | -3.09 | 0.002 | -22.04 | < 0.001 |
| High litter | -0.51 | 0.21 | -2.45 | 0.01 | | |
| *Microstegium* present | -0.21 | 0.22 | -0.97 | 0.33 | -1.57 | 0.21 |
| Low litter:*Microstegium* present | 0.06 | 0.28 | 0.21 | 0.83 | | |
| Medium litter:*Microstegium* present | 0.39 | 0.29 | 1.36 | 0.17 | -3.74 | 0.29 |
| High litter:*Microstegium* present | -0.10 | 0.29 | -0.34 | 0.74 | | |

**Table S4.** Model summary of *E. virginicus* disease incidence (proportion of tillers, binomial distribution, logit link). Estimates, standard errors (SE), z statistic, and associated *P*-value from maximal model (*n* = 48). Predictor variables without estimates were omitted during model selection (deviances and *P*-values from Chi-squared tests of nested models). Model comparison with continuous litter model: ΔAIC = 143.2.

| Predictor | Estimate | SE | z | Pr(>\|z\|) | Deviance | Pr(>Chi) |
|---|---|---|---|---|---|---|
| Intercept | -3.38 | 0.23 | -14.48 | < 0.001 | | |
| Low litter | 2.28 | 0.26 | 8.69 | < 0.001 | | |
| Medium litter | 1.02 | 0.28 | 3.61 | < 0.001 | -180.64 | < 0.001 |
| High litter | 1.67 | 0.26 | 6.31 | < 0.001 | | |
| *Microstegium* present | 2.14 | 0.27 | 7.82 | < 0.001 | -247.28 | < 0.001 |
| Low litter:*Microstegium* present | -0.79 | 0.33 | -2.37 | 0.02 | | |
| Med. litter:*Microstegium* present | -0.69 | 0.35 | -1.94 | 0.05 | -6.62 | 0.08 |
| High litter:*Microstegium* present | -0.44 | 0.33 | -1.32 | 0.19 | | |

**Table S5.** Model summary of log-transformed *M. vimineum* plant biomass. Estimates, standard errors (SE), t statistic, and associated *P*-value from maximal model (*n* = 48). Predictor variables without estimates were omitted during model selection (F statistics and *P*-values from comparisons of nested models). Model comparison with categorical litter model: ΔAIC = 2.1.

| Predictor | Estimate | SE | t | Pr(>\|t\|) | F | Pr(>F) |
|---|---|---|---|---|---|---|
| Intercept | -0.09 | 0.06 | -1.60 | 0.12 | | |
| Litter (g) | -0.05 | 0.03 | -1.63 | 0.11 | 5.90 | 0.02 |
| *Elymus* present | -0.10 | 0.08 | -1.26 | 0.22 | 4.36 | 0.04 |
| Litter (g):*Elymus* present | -0.004 | 0.04 | -0.10 | 0.92 | 0.01 | 0.92 |

**Table S6.** Model summary of log-transformed *E. virginicus* plant biomass. Estimates, standard errors (SE), t statistic, and associated *P*-value from maximal model ($n$ = 48). Predictor variables without estimates were omitted during model selection (F statistics and *P*-values from comparisons of nested models). Model comparison with continuous litter model: ΔAIC = 3.4.

| Predictor | Estimate | SE | t | Pr(>\|t\|) | F | Pr(>F) |
|---|---|---|---|---|---|---|
| Intercept | -1.88 | 0.14 | -13.05 | < 0.001 | | |
| Low litter | -0.10 | 0.20 | -0.50 | 0.62 | | |
| Medium litter | -0.002 | 0.20 | -0.01 | 0.99 | 2.20 | 0.10 |
| High litter | -0.03 | 0.20 | -0.16 | 0.88 | | |
| *Microstegium* present | -1.67 | 0.20 | -8.19 | < 0.001 | 321.3 | < 0.001 |
| Low litter:*Microstegium* present | -0.51 | 0.29 | -1.76 | 0.09 | | |
| Medium litter:*Microstegium* present | -0.38 | 0.29 | -1.33 | 0.19 | 1.33 | 0.28 |
| High litter:*Microstegium* present | -0.11 | 0.29 | -0.38 | 0.70 | | |

**Table S7.** Model summary of the residuals of *E. virginicus* establishment regressed against treatments (Table S3). Estimates, standard errors (SE), t statistic, and associated *P*-value are shown ($n$ = 48).

| Predictor | Estimate | SE | t | Pr(>\|t\|) |
|---|---|---|---|---|
| Intercept | -0.16 | 0.28 | -0.57 | 0.58 |
| Disease incidence | 0.61 | 0.84 | 0.73 | 0.47 |

**Table S8.** Model summary of the residuals of log-transformed *E. virginicus* plant biomass regressed against treatments (Table S6). Estimates, standard errors (SE), t statistic, and associated *P*-value are shown ($n$ = 48).

| Predictor | Estimate | SE | t | Pr(>\|t\|) |
|---|---|---|---|---|
| Intercept | 0.03 | 0.08 | 0.40 | 0.70 |
| Disease incidence | -0.14 | 0.23 | -0.59 | 0.56 |

**Table S9.** Model summary of the relative abundance of *M. vimineum* (log(*M. vimineum* biomass/total biomass)). Estimates, standard errors (SE), t statistic, and associated *P*-value from maximal model (*n* = 24). F statistic and *P*-value from comparison of nested models. Model comparison with continuous litter model: ΔAIC = 7.8.

| Predictor | Estimate | SE | t | Pr(>\|t\|) | F | Pr(>F) |
|---|---|---|---|---|---|---|
| Intercept | -0.036 | 0.004 | -9.99 | < 0.001 | | |
| Low litter | 0.018 | 0.005 | 3.55 | 0.002 | | |
| Medium litter | 0.008 | 0.005 | 1.57 | 0.13 | 4.28 | 0.02 |
| High litter | 0.007 | 0.005 | 1.35 | 0.19 | | |
