## Appendix S3 for "Invasive grass litter suppresses native species and promotes disease"

**Appendix S3** for "Invasive grass litter suppresses native species and promotes disease " by Liliana Benitez, Amy E. Kendig, Ashish Adhikari, Keith Clay, Philip F. Harmon, Robert D. Holt, Erica M. Goss, and S. Luke Flory

**Overview**

We performed a greenhouse experiment to test the effects of moisture on litter-induced disease of *Microstegium vimineum* and to assess the potential for *Microstegium* litter to promote disease symptoms on five native species: *Calamagrostis canadensis*, *Dichanthelium clandestinum*, *Elymus virginicus*, *Eragrostis trichodes*, and *Glyceria striata*. We refer to each species by its genus for the remainder of the Appendix. We hypothesized that moisture, and leaf wetness in particular, would increase disease incidence on *Microstegium* (Green et al. 2004). We hypothesized that litter would increase disease symptoms on *Elymus* and *Glyceria*, which are susceptible to *Bipolaris* associated with *Microstegium* (Flory et al. 2011, Lane et al. 2020). We were unsure how litter would affect disease symptoms on the other three native species.

**Methods**

We purchased seeds of *Calamagrostis*, *Elymus*, *Eragrostis*, and *Glyceria* from Prairie Moon Nursery (Winona, MN, USA) in spring 2018 and we purchased *Dichanthelium* seeds from Sheffield's Seed Company (Locke, NY, USA) in spring 2018. The *Microstegium* seeds and litter were collected from Big Oaks National Wildlife Refuge (Madison, IN, USA), as described in the main text. Additional litter was collected from the same field site in June 2018 and homogenized with the litter collected in May 2018. In contrast with the experiment described in the main text, we did not remove non-*Microstegium* litter or cleistogamous *Microstegium* seeds from the litter. Seeds and litter were stored at 4°C until the start of the experiment.

The experimental design included five replicate pots for each treatment. There were three treatments for *Microstegium*—no litter, litter with low moisture, and litter with high moisture—and two treatments for the native species—no litter and litter with high moisture. We planted each species in its own pot (1 L, 6 in. diameter) using Metromix 360 growing medium (Sungro Horticulture, Agawam, MA, USA) saturated with tap water. We used data from a pilot experiment to determine how many seeds to add for each species and when to plant them so that they would be approximately the same height and density when litter was added to the pots. This goal was motivated by the assumption that plant height and density affect the distance fungal spores must travel from litter to live plants for disease transmission to occur. We planted 60 *Glyceria* seeds per pot 24 days prior to litter addition. We planted 1000 *Calamagrostis* seeds, 80 *Dichanthelium* seeds, 60 *Elymus* seeds, 80 *Eragrostis* seeds, and 40 *Microstegium* seeds per pot 20 days prior to litter addition. Approximately seven days post planting, we thinned *Elymus* and *Microstegium* pots to 20–25 individuals each. *Calamagrostis* and *Glyceria* pots were thinned to space out individuals and the other two native species were not thinned.

On July 26, 2018, we filled one-gallon plastic bags with 9.5 g of *Microstegium* litter and three paper towels saturated with deionized water (one bag per experimental pot). We stored the bags in a growth chamber at 24°C and a 16/8-hour light/dark cycle for 89.5 hours prior to being added to experiment pots. Bags were checked daily and water was added as needed. We measured the heights of five plants per pot from the soil surface in the 24 hours leading up to litter addition. We attached 13 in. x 20 in. clear plastic cylindrical humidity chambers (0.005 Grafix Dura-Lar® film, Maple Heights, OH) around each pot. Just prior to adding litter to pots with the high moisture treatment, we misted plants with tap water for five seconds. We poured the litter from each plastic bag onto a pot, added tap water to the empty bag, shook it, and poured it on the pot. For the high moisture treatment, we misted plants with tap water for five seconds after litter had been added to all pots and two more times afterwards at two-hour intervals. High moisture pots were then misted every two hours and low moisture pots were misted once per day for the second and third days from 9:30 AM to 1:30 PM. We recorded whether water droplets were observed on leaf surfaces in the litter addition treatments ten times over the first 45 hours of the experiment. In the first 19.4 hours, pots with any water droplets were recorded, but after this time, pots were only recorded as having water droplets if droplets were observed on at least three leaves.

On September 13, 2018, plant stems were clipped at the soil surface with scissors and stored at 4°C in paper bags. Over the following month, we examined plants for foliar lesions for one minute per bag or until lesions were found. Pots were considered infected if at least two brown lesions were observed on leaf surfaces. Black spots were observed but were not counted as lesions. Because *Microstegium* germinated from cleistogamous seeds in the litter, we also noted whether lesions were observed on their leaf surfaces.

We used Fisher's Exact Test to evaluate whether adding litter to pots significantly increased the number of pots with lesions and whether the high moisture treatment significantly increased the number of *Microstegium* pots with lesions compared to the low moisture treatment. Analyses were performed in R version 4.0.1 (R Core Team 2020).

**Results and Discussion**

The average height of plant species at the time of inoculation ranged from 2.43 cm (*Glyceria*) to 12.2 cm (*Elymus*) (Fig. 1), suggesting that fungal spores may have more easily travelled from litter to live plants for *Glyceria* than *Elymus*.

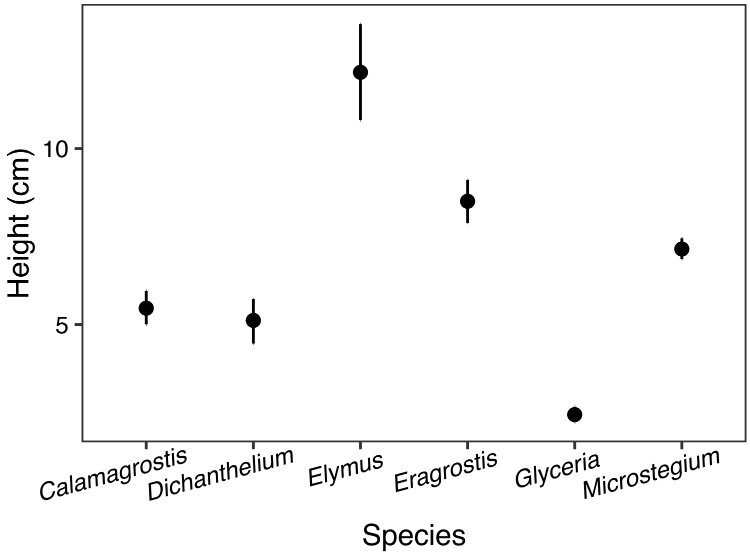


**Figure 1.** The average height of 25 plants (five plants per pot) for each of the species at the time of litter addition (mean ± 95% CI). We aimed to have plants at a similar height so that the distance from that fungal spores travelled from litter to live plants was consistent across the species.

Plants in one *Glyceria* pot and two *Calamagrostis* pots were completely covered by litter following the first 45 hours of litter addition and could not be assessed for leaf wetness. For the rest of the pots, leaves were consistently wet following litter addition except for *Microstegium* with the low moisture treatment. Therefore, the moisture treatments affected leaf wetness as intended.

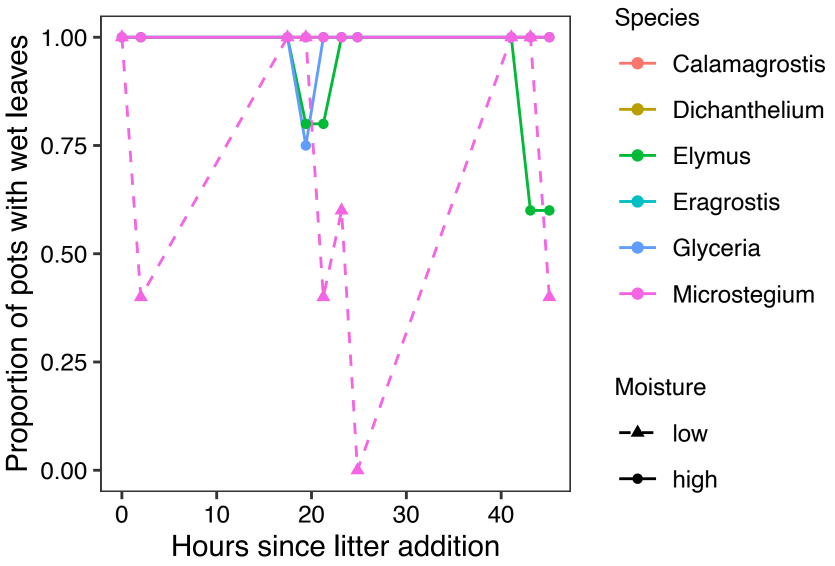


**Figure 2.** The low moisture treatment reduced the proportion of *Microstegium* pots with wet leaves. All other pots, which received the high moisture treatment, had consistently wet leaves.

The high moisture treatment significantly increased the number of *Microstegium* pots with foliar lesions compared to the low moisture treatment (odds ratio = 0, 95% CI = 0–0.98, $P$ = 0.047, Fig 3a). This result suggests that foliar fungal lesions are more likely to form on *Microstegium* leaves when they are wet, consistent with a previous study demonstrating that longer leaf wetness duration increased disease severity by *Bipolaris gigantea* on green foxtail (Green et al. 2004).

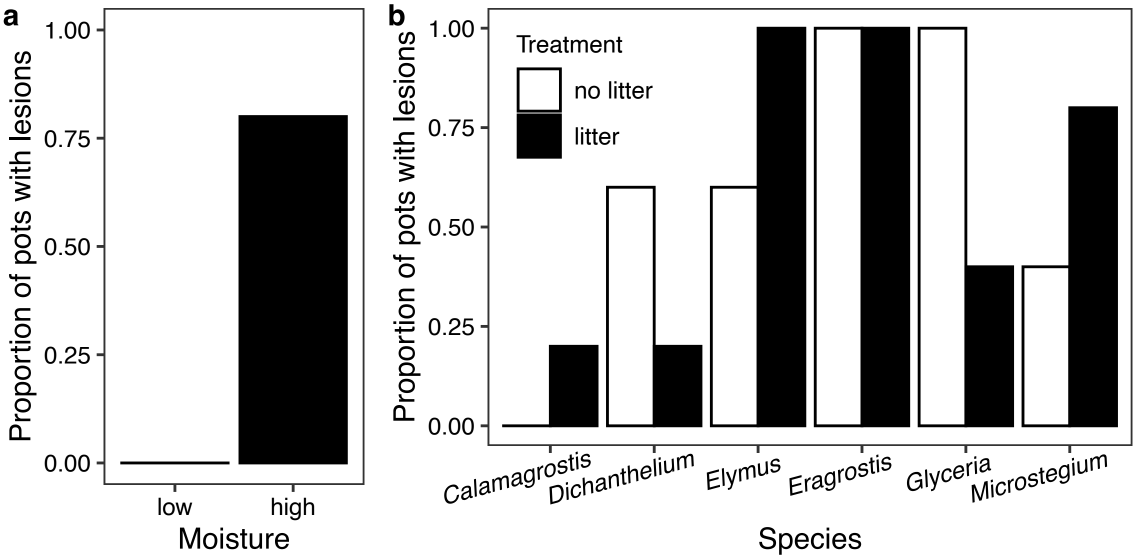


**Figure 3.** The high moisture treatment, which increased leaf wetness, significantly increased the proportion of *Microstegium* pots with foliar lesions (b) but adding litter did not significantly increase the proportion of pots with foliar lesions for any of the species (a).

Adding litter to pots did not significantly increase the proportion of pots with lesions for any of the species (Table 1). However, *Calamagrostis*, *Elymus*, and *Microstegium* tended to have more pots with lesions when litter was added, suggesting that litter promoted foliar fungal infection on these three species (Fig. 3b). Multiple species had lesions even without litter, suggesting alternative sources of fungal infection in the greenhouse, such as contaminated soil. Adding litter tended to decrease the proportion of pots with lesions for *Dichanthelium* and *Glyceria*. Therefore, litter may have blocked external sources of fungi from contacting *Dichanthelium* and *Glyceria* leaf surfaces. This result was particularly surprising for *Glyceria* because its leaves were close to the litter (Fig. 1) and it is susceptible to *Bipolaris* fungi associated with *Microstegium* (Flory et al. 2011). Other sources of infection may have driven lesion patterns observed on Glyceria, overshadowing *Bipolaris* transmission from litter.

**Table 1.** Fisher's exact tests for the number of pots with foliar lesions with and without litter addition.

| Species | Odds ratio | 95% CI | *P* |
|---|---|---|---|
| *Calamagrostis* | 0.00 | 0.00–39.00 | 1.00 |
| *Dichanthelium* | 4.9 | 0.21–390.56 | 0.52 |
| *Elymus* | 0.00 | 0.00–5.12 | 0.44 |
| *Eragrostis* | 0.00 | 0.00–infinity | 1.00 |
| *Glyceria* | infinity | 0.49–infinity | 0.17 |
| *Microstegium* | 0.20 | 0.003–4.59 | 0.52 |

R Core Team. 2020. R: A Language and Environment for Statistical Computing. R Foundation for Statistical Computing, Vienna, Austria.
